# Using *SAFEPATH* to Uncover a Novel Mechanism of Hepatotoxicity for Gefitinib

**DOI:** 10.1101/2024.07.31.605988

**Authors:** Layla Hosseini-Gerami, Sara Masarone, Jordan Lane

## Abstract

This white paper details the research conducted by Ignota Labs using their advanced causal and explainable AI technology, ***SAFEPATH***, to analyse the mechanisms of hepatotoxicity for two EGFR-TKI inhibitors, Erlotinib and Gefitinib, the latter having an as yet unknown mechanism of toxicity. The known mechanism of UGT1A1-mediated toxicity of Erlotinib was recovered, and a novel sphingolipid metabolism mechansim of toxicity of Gefitinib was hypothesised and subsequently experimentally validated. Crucially, we were also able to suggest the reason for the observed heterogeneous toxicity response to Gefitinib. This study exemplifies the potential of integrating AI tools with comprehensive datasets to improve drug safety and patient management.

## 1 Introduction

Erlotinib and Gefitinib, both efficacious treatments for non-small cell lung cancer (NSCLC), are classified by the FDA as “Most-DILI-Concern” [1] due to their severe hepatotoxic effects. Despite their clinical success, a considerable proportion of patients experience liver toxicity, necessitating frequent monitoring and dose adjustments [2, 3].

Liver toxicity associated with these drugs poses significant challenges in clinical settings, impacting patient compliance and treatment outcomes. Understanding the mechanisms behind this toxicity is crucial for improving patient management and developing safer therapeutic alternatives. This paper presents Ignota Labs’ innovative approach to uncovering the underlying mechanisms of this toxicity, which remained elusive for over two decades.

## 2 Methodology

Ignota Labs leveraged the DILIrank dataset, comprising 1,036 FDA-approved drugs categorized by their liver injury risk. The DILIrank dataset includes three categories: Most-DILI-Concern, Less-DILI-Concern, and No-DILI-Concern, based on verified causal links between a medication and liver damage.

Using the proprietary AI platform, ***SAFEPATH***, Ignota Labs mapped potential pathways and validated their predictions through wet-lab experiments. The focus was on identifying novel pathways and validating known ones to provide a comprehensive understanding of the hepatotoxicity mechanisms for Erlotinib and Gefitinib. The AI platform integrates various data types, including transcriptomic, bioactivity, and pathway data, to build a robust understanding of the mechanism of Drug Induced Liver Injury (DILI). The platform then carries out a causal analysis of pathways leading to toxicity, giving an interpretable explanation of complex clinical outcomes.

### 2.1 Building a Comprehensive Knowledge Graph for DILI

***SAFEPATH*** leverages a knowledge graph (KG) of heterogeneous biological relationships to make connections between drugs and toxicity. As shown in Table 1, the different types of relationships in our knowledge graph (DILI-KG) serve various purposes.

**Table 1:** Types of Relationships in the Drug Induced Liver Injury Knowledge Graph (DILI-KG) and Their Purposes.

| Relationship Type | Purpose |
| --- | --- |
| Drug-Protein | To understand on- and off-target effects, including metabolic enzymes. Enriched with predicted edges using our target prediction tool **CELLSCAPE** trained using our state-of-the-art IgnotaNet architecture. |
| Drug-Gene | To provide information on changes in gene expression induced by drug treatment. |
| Protein-Protein | To understand protein signalling patterns and connections. |
| Protein-Pathway | To connect proteins to the biological processes they participate in. |
| Drug-Adverse Event | To provide details of unwanted effects manifesting from drug treatment. This includes our proprietary 60,000+ compound mitochondrial toxicity and cytotoxicity dataset. |

The full DILIrank dataset (1,036 compounds) was supplemented with additional data using our in-house data processing pipeline to form a set of 1,888 compounds. The final DILI dataset comprised of 1,033 toxic and 855 non-toxic compounds.

Ignota Labs have developed a KG generation pipeline, based on a set of drugs under investigation (here, our DILI dataset). This pipeline traverses public and proprietary datasets and predictions, standardises the information, and integrates it into a causal structure. A toy example of a path in the DILI-KG is shown in Figure 1, where drug *inhibits* protein, which *participates in* a pathway, which *causes* toxicity. We also know that drug *causes* toxicity.

**Figure 1:**
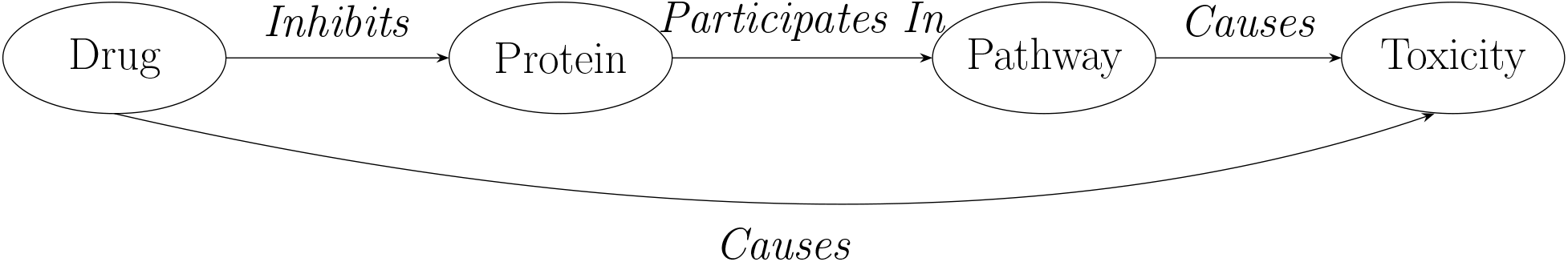
Toy example of a path in the DILI-KG.

### 2.2 Training an Explainable DILI AI Model

***SAFEPATH*** ‘s Causal AI architecture was leveraged with our DILI-KG to train a model to predict DILI. The 1,888 drugs were split in a class-stratified fashion into train-validation-test with a ratio of 0.7-0.2-0.1 where Erlotinib and Gefitinib were placed in the test set. Feature selection using a method based on the Relief algorithm [4] was applied to the training set only to select the top *n* protein targets for inferring DILI, where *n* is a hyperparameter optimised during tuning. The validation set was used to tune this parameter as well as other parameters relating to the types and amount of data extracted from the DILI-KG using a Bayesian optimisation method.

The best performing model based on the validation set included 400 off-target predictions and their selected relationships in the DILI-KG. This was applied to the held-out test set and chosen as our final model. Although the focus here is on understanding rather than predicting, the high performance of the model (validation F1=0.74, test F1=0.73) shows that the model is capable of learning about the different relationships in the KG that contribute towards DILI.

### 2.3 Transcriptomic Validation Experiments

Primary rat hepatocytes were treated with Gefitinib for 16h. The mRNA was extracted and analysed with the Affymetrix whole-genome Rat GeneChip 230 2.0 Array. The data were quality controlled, pre-processed and used with statistical analyses to compute differential gene expression between Gefitinib and DMSO vehicle control. The resulting log2-Fold Change and Benjamini-Hochberg corrected p-values were used as input to GSEA [5] to compute pathway enrichment on the transcriptional level.

We then applied our transcriptional analysis platform, ***GENESCAPE*** to generate a protein-protein interaction (PPI) subnetwork connecting targets to modulated transcription factors. This method elucidates the specific protein signalling changes induced after target engagement by taking into account the ground-truth transcriptional data.

### 2.4 Kinase Enzyme Assay Validation Experiments

Erlotinib and Gefitinib were profiled against Protein Kinase D1 via radiometric HotSpot™ kinase assay at 10 different concentrations, using Staurosporine as positive control and DMSO as vehicle. Resulting data were analysed with GraphPad to produce IC50 curves.

## 3 Results

### 3.1 *SAFEPATH* Elucidates Known and Novel Mechanisms of Toxicity

The output from ***SAFEPATH*** ‘s analysis has been summarised in Figure 2, and can be split into three broad buckets:

**Figure 2:**
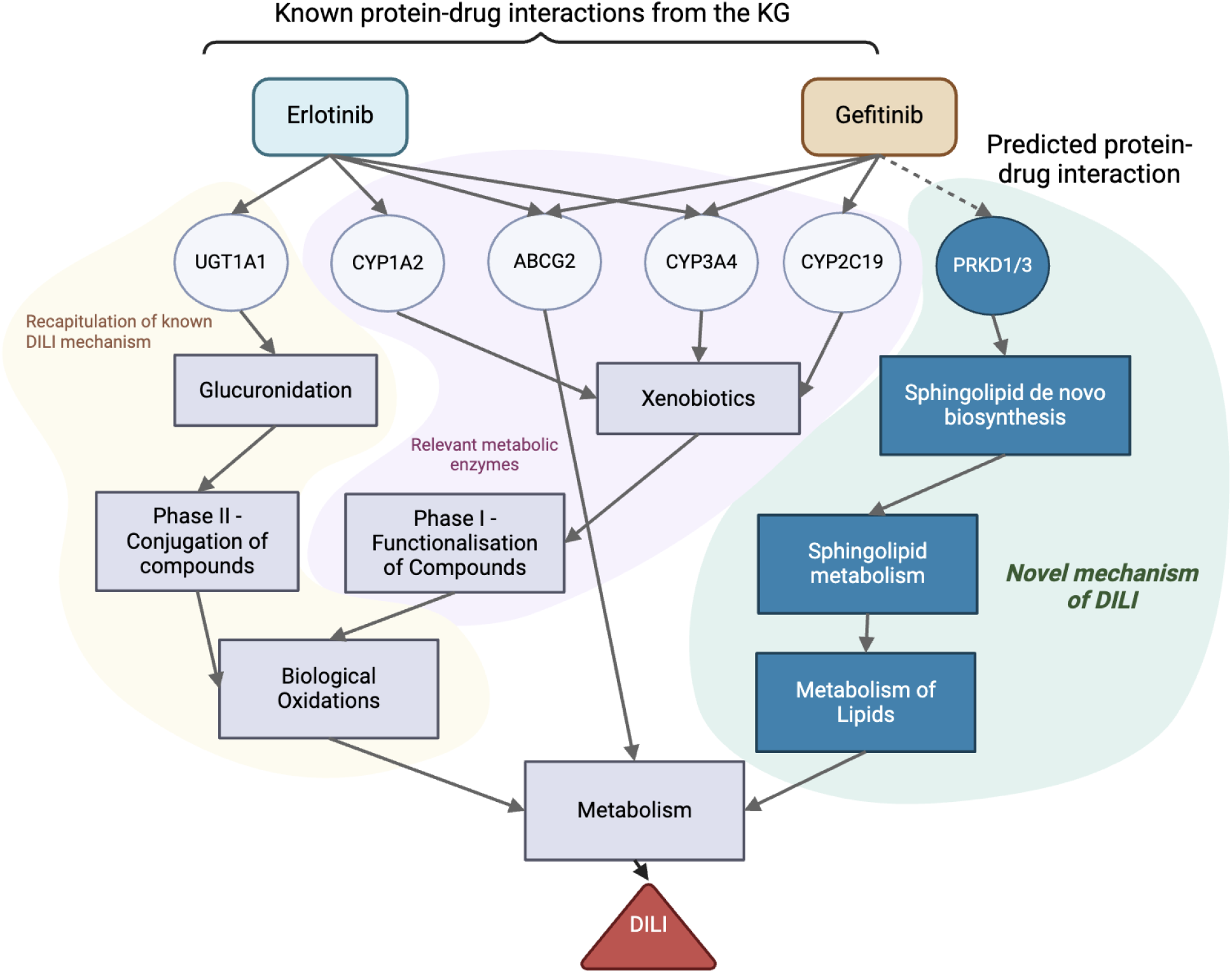
Output of *SAFEPATH*.

- Known Erlotinib UGT1A1 Mechanism (left)
- Known Metabolism Enzymes (middle)
- Novel Gefitinib Mechanism (right)

#### 3.1.1 Known Erlotinib UGT1A1 Mechanism

UGT1A1 is a uridine diphosphate(UDP)-glucuronosyltransferase which carries out the process of glucorondiation, or the transfer of glucuronic acid, on its substrates [6] Erlotinib is a potent inhibitor of UGT1A1 (sub-*µ*M) which is known to increase the risk of drug–drug interactions (DDIs), hyperbilirubinemia and DILI [7]. Aggressive cancers are often treated with multiple drugs, and the consequences of this are exemplified with the interaction between Erlotinib and Irinotecan which cases severe toxicity, as the hepatotoxic metabolite of Irinotecan, SN-38, cannot be deactivated by UGT1A1 after Erlotinib inhibition [8] (Figure 3).

**Figure 3:**
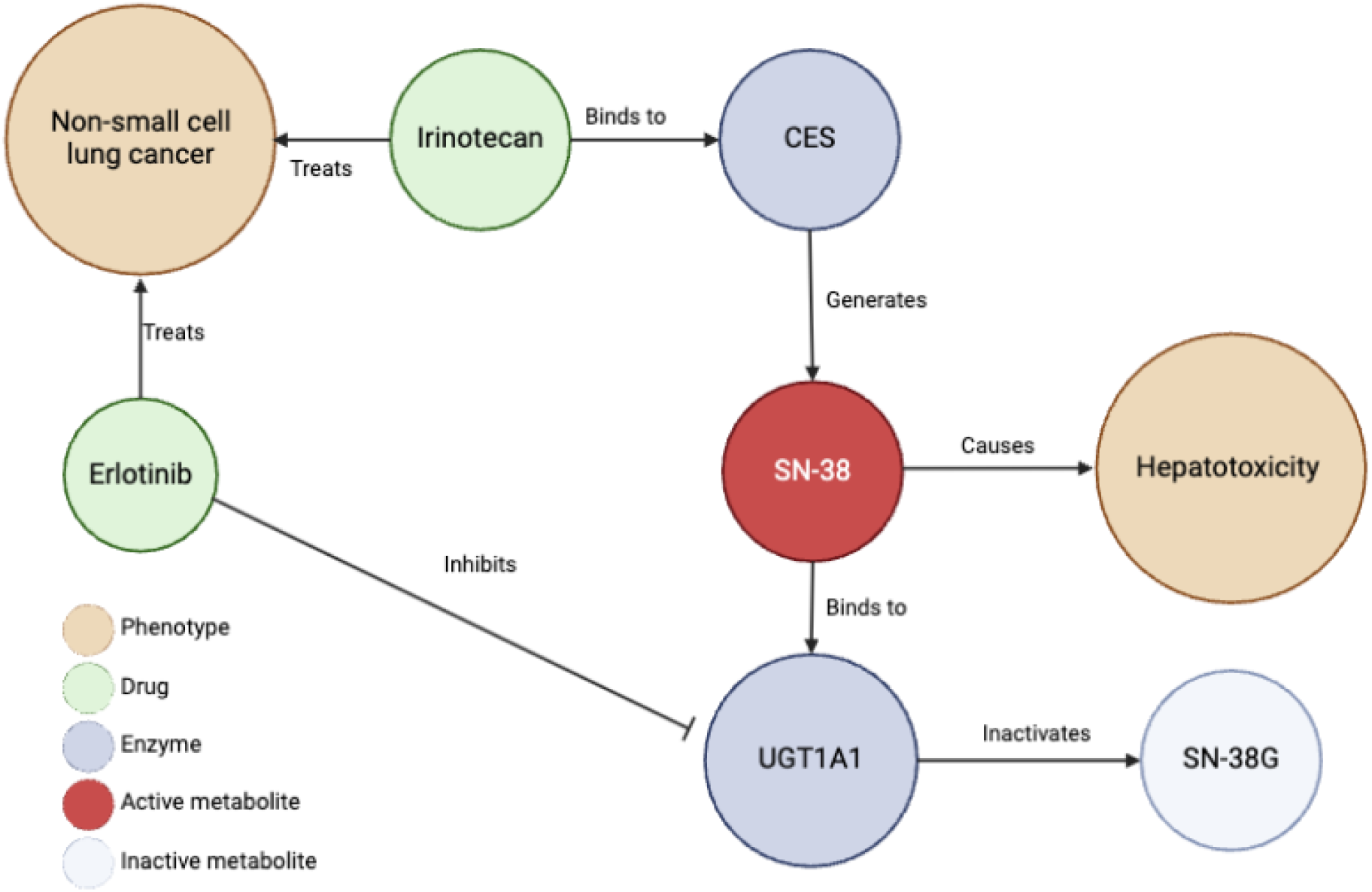
Inhibition of UGT1A1 by Erlotinib leads to an accumulation of Irinotecan’s active metabolite SN-38, as it cannot be deactivated. This effect is exacerbated further in less active UTG1A1*28 allele which is commonly found in African-Americans (0.42–0.45 allele frequency) and Caucasians (0.26–0.31), and is less common in Asian populations (0.09–0.16). *CES - carboxylesterase, UGT1A1-UDP-glucuronosyltransferase 1, SN-38 - active metabolite of irinotecan, SN-38G-inactive metabolite of irinotecan*.

#### 3.1.2 Known Metabolism Enzymes

Known metabolic enzymes were uncovered, which contribute towards DILI. These include cytochrome P450 (CYP) enzymes and ABCG2, which have been implicated in Erlotinib-induced DILI [9].

#### 3.1.3 Novel Gefitinib Mechanism

A novel pathway of toxicity for Gefitinib was revealed *via* Protein Kinase D1 and Protein Kinase D3 (PRKD1 and PRKD3), enzymes involved in sphingolipid metabolism. Disruption in sphingolipid homeostasis is linked to hepatocellular death and liver injury [10], however the link between this mechanism and Gefitinib had not yet been established.

This link was predicted by ***CELLSCAPE***, which predicted that Gefitinib, but not Erlotinib, inhibits PRKD1 and PRKD3 at concentrations above 10 µM (Table 2). This prediction was based solely on the chemical structures of the compounds.

**Table 2:**
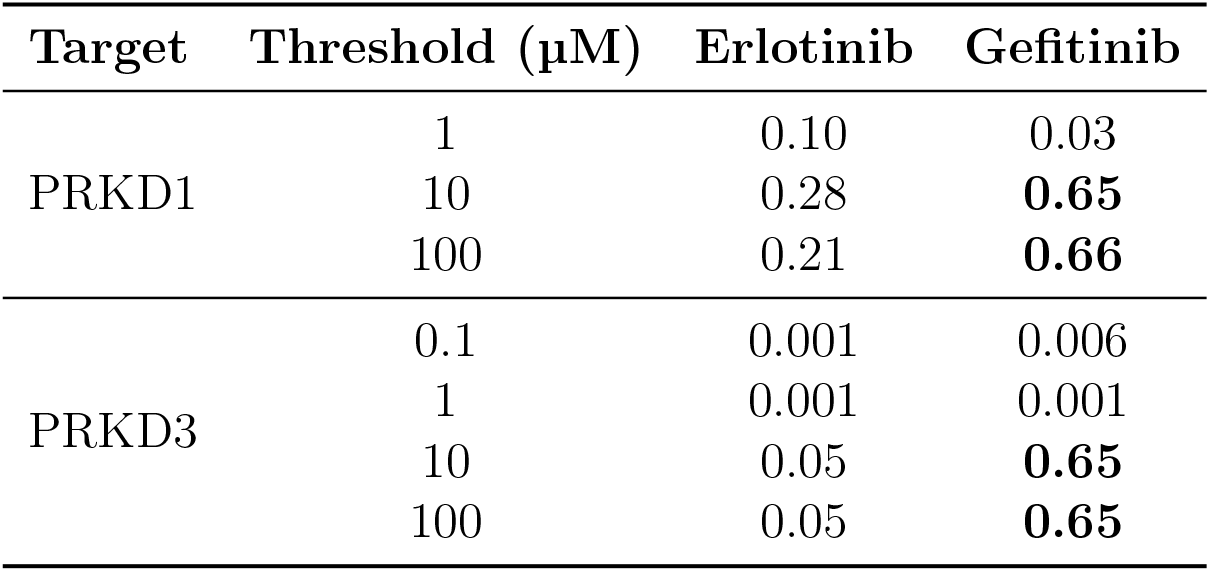
Predicted probability of target engagement at different concentrations (µM) of Erlotinib and Gefitinib for PRKD1 and PRKD3 targets. Bold values highlight significant thresholds (0.5).

| Target | Threshold ($\mu\text{M}$) | Erlotinib | Gefitinib |
| --- | --- | --- | --- |
| PRKD1 | 1 | 0.10 | 0.03 |
|  | 10 | 0.28 | **0.65** |
|  | 100 | 0.21 | **0.66** |
| PRKD3 | 0.1 | 0.001 | 0.006 |
|  | 1 | 0.001 | 0.001 |
|  | 10 | 0.05 | **0.65** |
|  | 100 | 0.05 | **0.65** |

### 3.2 Validation of Novel Mechanism

#### 3.2.1 Validation of Gefitinib Sphingolipid Metabolism Pathway

Transcriptomic analysis of Gefitinib-treated primary rat hepatocytes revealed enrichment of sphingolipid metabolism pathways post-treatment (Table 3). The data showed significant downregulation of genes related to sphingolipid homeostasis, validating the AI-predicted pathway.

**Table 3:**
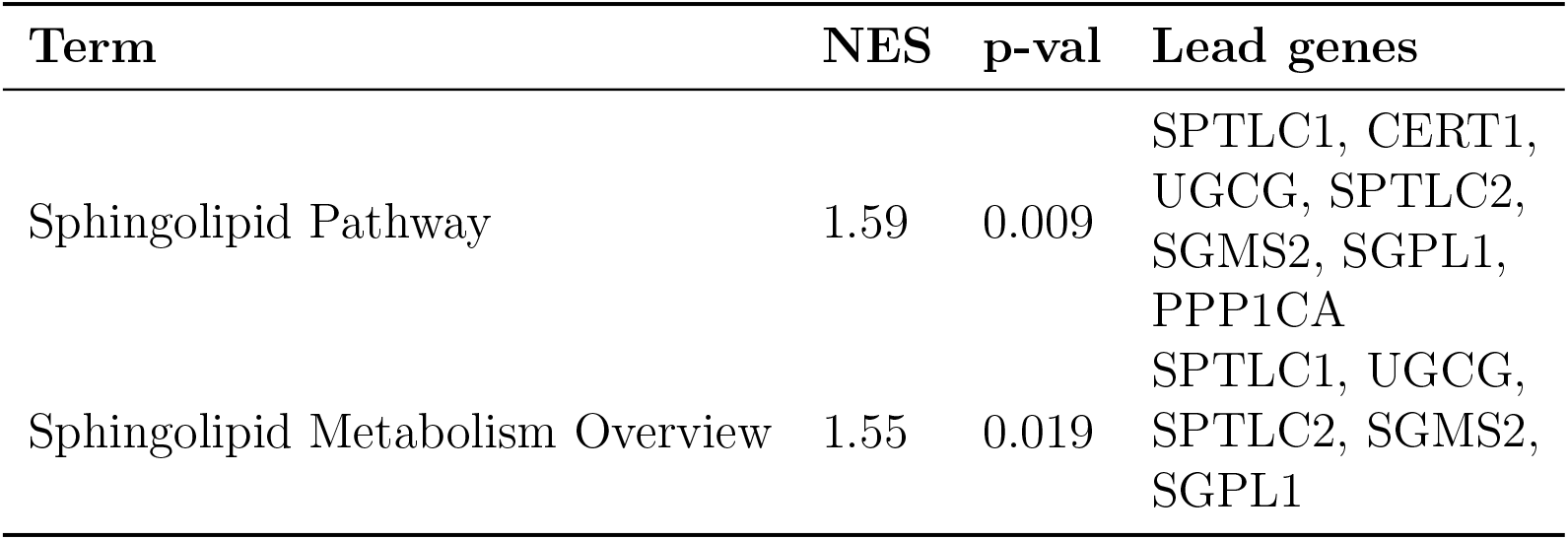
Normalised enrichment score (NES) and p-value for Sphingolipid Pathway and Sphingolipid Metabolism Overview with their respective lead genes altered by Gefitinib, based on GSEA analysis of rat hepatocyte mRNA data.

| Term | NES | p-val | Lead genes |
| --- | --- | --- | --- |
| Sphingolipid Pathway | 1.59 | 0.009 | SPTLC1, CERT1, UGCG, SPTLC2, SGMS2, SGPL1, PPP1CA |
| Sphingolipid Metabolism Overview | 1.55 | 0.019 | SPTLC1, UGCG, SPTLC2, SGMS2, SGPL1 |

#### 3.2.2 Validation of PRKD1 inhibition

Wet-lab validations confirmed Gefitinib’s binding to PRKD1 and lack of activity of Erlotinib (Figure 4). The binding assays demonstrated that Gefitinib inhibits PRKD1 at 3 µM, which is close to its maximum concentration (cMax) of 2.5-5 µM observed in patients at a dose of 700-1000 mg/d [11].

**Figure 4:**
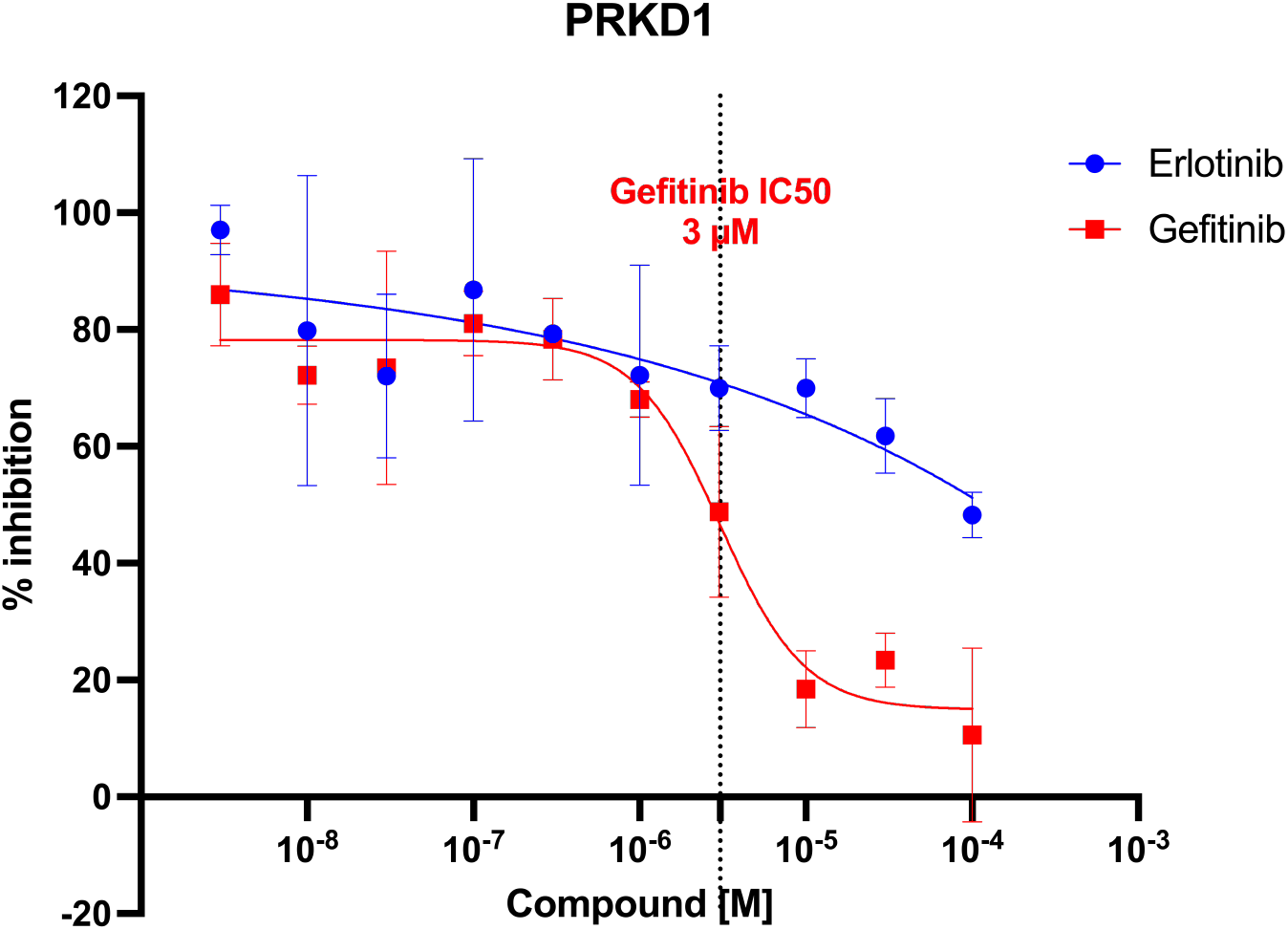
Kinase Enzyme Assay experiments revealed that Gefitinib inhibits PRKD1 at 3 µM.

Two other tyrosine kinase inhibitors, Sunitinib and Crizotinib, are also inhibitors of PRKD1 (Kd 310nM and 990nM, respectively [12]) and are known to cause hepatotoxicity [13, 14]. This further strengthens the causative link between PRKD1 inhibition and DILI.

#### 3.2.3 Validation of Mechanistic Cascade

***GENESCAPE***, our transcriptional analysis platform, used known protein-protein interactions to connect PRKD1 inhibition to Gefitinib-induced transcriptional changes. This revealed a known link between PRKD1 and the modulation of Sphingosine Kinase 2 (SPHK2) *via* SYK, PRKCD and MAPK1, which is concordant with the observed transcriptional changes (Figure 5). This signalling cascade is present in the sphingolipid signalling pathway, where the crucial MAPK1/SPHK interaction leads to the transformation of Sph (Sphingosine) to S1P (Sphingosine 1-phosphate) [15].

**Figure 5:**
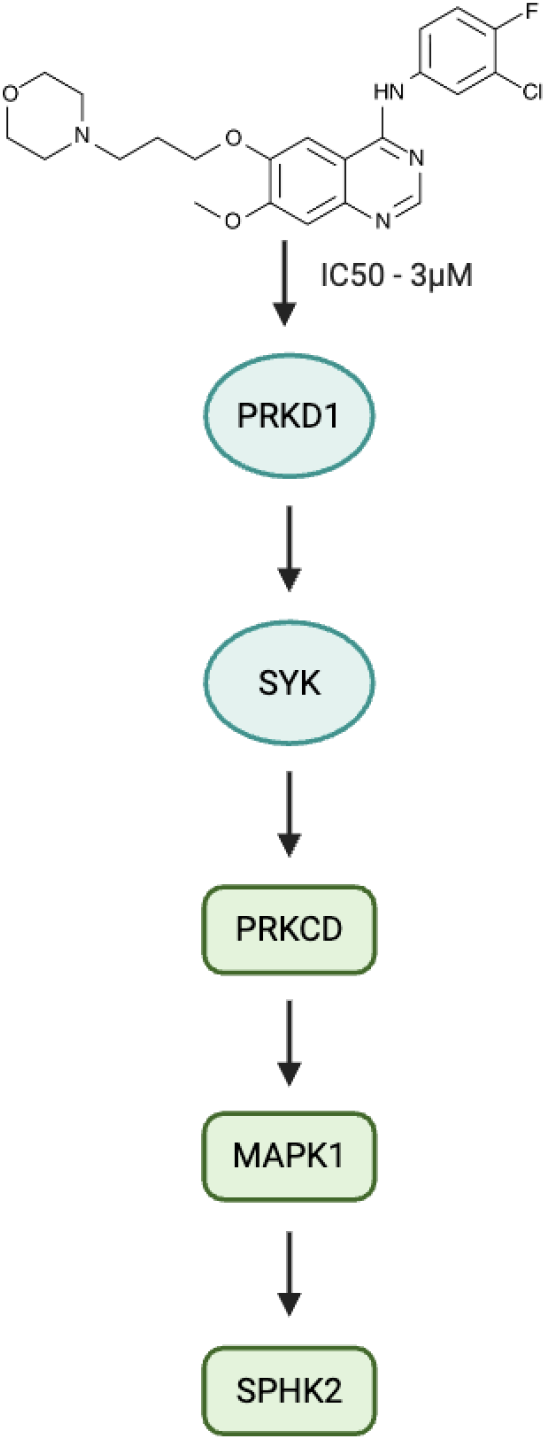
Highlighted pathway on the ***GENESCAPE*** subnetwork showing how PRKD1 modulation leads to changes in SPHK2 *via* SYK, PRKCD and MAPK1.

## 4 Clinical Relevance

The findings offer crucial insights into managing NSCLC patients on Gefitinib. Understanding this novel mechanism can guide personalised dosing strategies, reducing the risk of liver damage. Moreover, monitoring patients’ drug pharmacokinetics and maximum concentration (cMax) can help tailor treatment to individual metabolism profiles without compromising therapeutic efficacy.

The validated pathways for Erlotinib and Gefitinib provide a blueprint for investigating other drugs with unresolved toxicity issues. These findings highlight the importance of integrating advanced AI tools and comprehensive datasets to enhance understanding of drug toxicity mechanisms and improve patient management strategies.

For Erlotinib, the known mechanism of UGT1A1 inhibition underscores the need for routine liver function monitoring, especially in patients with genetic predispositions to reduced UGT1A1 activity. For Gefitinib, the novel mechanism involving PRKD1 and sphingolipid metabolism suggests new avenues for therapeutic interventions and safer drug design.

## 5 Implications for Future Research

This study demonstrates the transformative potential of combining AI with comprehensive datasets to unravel complex drug toxicity mechanisms. Future research can focus on refining these AI models and exploring additional datasets to further enhance drug safety profiles. The validated mechanisms for Erlotinib and Gefitinib provide a strong foundation for developing predictive models for other drugs, potentially reducing the incidence of drug-induced liver injury.

Additionally, the insights gained from this research can inform the design of new drugs that retain therapeutic efficacy while minimising hepatotoxic risks. By understanding the molecular drivers of toxicity, pharmaceutical companies can develop targeted interventions to mitigate these effects.

## 6 Conclusion

Ignota Labs’ research marks a significant advancement in understanding the hepatotoxic mechanisms of Erlotinib and Gefitinib. The integration of advanced AI tools and thorough experimental validation underscores the potential of such approaches in drug safety research. These findings pave the way for improved patient management and the development of safer therapeutic alternatives, highlighting the transformative impact of AI in pharmaceutical research.

